# Hydrogel injection molded complex macroencapsulation device geometry improves long-term cell therapy viability and function in the rat omentum transplant site

**DOI:** 10.1101/2024.08.11.607495

**Authors:** Amy E. Emerson, Quincy Lyons, Matthew W. Becker, Keven Sepulveda, Shivani Hiremath, Sarah R. Brady, Jessica D. Weaver

## Abstract

Insulin-secreting allogeneic cell therapies are a promising treatment for type 1 diabetes, with the potential to eliminate hypoglycemia and long-term complications of the disease. However, chronic systemic immunosuppression is necessary to prevent graft rejection, and the acute risks associated with immunosuppression limit the number of patients who can be treated with allogeneic cell therapies. Islet macroencapsulation in a hydrogel biomaterial is one proposed method to reduce or eliminate immune suppression; however, macroencapsulation devices suffer from poor oxygen transport and limited efficacy as they scale to large animal model preclinical studies and clinical trials. Hydrogel geometric device designs that optimize nutrient transport combined with methods to promote localized vasculogenesis may improve in vivo macroencapsulated cell viability and function. Here, we demonstrate with finite element modeling that a high surface area-to-volume ratio spiral geometry can increase macroencapsulated islet viability and function relative to a traditional cylindrical design, and we validate these observations *in vitro* under normoxic and physiological oxygen conditions. Finally, we evaluate macroencapsulated syngeneic islet survival and function in vivo in a diabetic rat omentum transplant model, and demonstrate that high surface area-to-volume hydrogel device designs improved macroencapsulated syngeneic islet function relative to traditional device designs.

## 1. Introduction

Type 1 Diabetes (T1D) is an autoimmune disease that systematically destroys the insulin-producing β cells in pancreatic islets^1^. Although T1D can be managed through blood glucose monitoring and exogenous insulin administration, imperfect management and inevitable blood glucose fluctuations result in an increased risk of long-term complications such as heart disease^2^, kidney disease^3^, and nerve damage^4^ which results in reduced quality of life and greater mortality. Islet transplantation is a promising cell therapy for T1D with the potential to eliminate blood glucose fluctuations and resultant long-term complications, but its widespread application is hindered by the need for chronic systemic immunosuppression^5^. Systemic immunosuppression carries the risk of adverse effects such as nephrotoxicity^6^, infections^7^, and cancer^8^, the acute risks of which typically outweigh the benefits of islet transplantation for all but the sickest patients (e.g. kidney failure or frequent life-threatening hypoglycemic events). Additionally, immunosuppressive drugs negatively affect the functionality of transplanted islets^9^, and immunosuppression may lead to increased insulin resistance^10^ resulting in ineffectual transplanted islets. As such, elimination or reduction of chronic systemic immunosuppression is critical to the translation of cell therapies for the treatment of T1D.

Pancreatic islet encapsulation within a hydrogel biomaterial may reduce or eliminate the need for chronic immunosuppression in T1D patients by preventing contact-mediated direct antigen recognition by the recipient’s immune system^11^. Alginate microencapsulation has been explored for the past 50 years^12,13^ with substantial preclinical success, but clinical success has been limited by multiple challenges, including insufficient oxygen transport to encapsulated cells, poor graft retrievability, and device fibrosis^14^. A curative dose in human microencapsulated islet transplantation requires a site large enough to accommodate hundreds of throusands of microcapsules ^15^, and capsule transplantation has been historically limited to the abdominal cavity, where capsules become fibrosed and adhere to abdominal tissues^16^.

Macroencapsulation is an alternative encapsulation strategy that prioritizes safety and retrievability by containing large numbers of cells within a single device. This approach is being pursued commercially, and several preclinical^17^ and clinical trials^18,19^ have demonstrated the potential of macroencapsulation devices in islet transplantation. One major drawback of existing macroencapsulation device designs is insufficient oxygen and nutrient transport to encapsulated cells, as tissue oxygen concentration drops precipitously 150-200 µm from vasculature ^20^. Cell therapy company Viacyte recently published data from a Phase I/II trial of non-immunoisolating macroencapsulation devices in a traditional cylindrical geometry at a high cell loading of ∼167 IEQ/µL, and histological assessment demonstrated large areas devoid of encapsulated cells, suggesting graft loss due to poor oxygen transport^21,22^. Insufficient oxygen and nutrients is particularly challenging for highly metabolic islets, where tissue oxygen saturation levels less than 3%^23^ result in loss of islet function, and below 1% results in hypoxia-induced islet death ^24^. Some strategies for supplying oxygen to islets in macroencapsulation devices are accelerating vascularization^25^, creating an oxygen supply within the implant^26^, or shortening diffusion distances within the macroencapsulation device^27^ by targeting geometries with an increased surface-area-to-volume (SA:V) ratio ^28–30^. Strategies that accelerate vascularization have demonstrated improvements in encapsulated islet survival and function, particularly in highly vascularized transplant sites like the omentum ^31–34^, but this technique has not yet been combined with high SA:V macroencapsulation gemoetries.

Macroencapsulation devices with complex, high SA:V geometries can be generated rapidly and reproducibly using hydrogel injection molding ^29,30^. Hydrogel injection molding is high-throughput, compatible with a range of natural and synthetic hydrogels, and is amenable to scaling in automated biomanufacturing ^29,30^. In this work, we designed high SA:V spiral geometry alginate macroencapsulation devices to test whether improved oxygen transport can enhance encapsulated islet viability and function *in vitro* and *in vivo* in a rat omentum transplant model (**Figure 1A**). We found through computational modeling of oxygen transport that high SA:V spiral encapsulation devices increased the functional islet loading density relative to standard geometry lower SA:V devices at physiological oxygen levels. We optimized our slow-gelling alginate formulation to hydrogel injection mold the high SA:V spiral design, and validated our computational models with pseudoislets and primary rat islets. Finally, we evaluated the impact of macroencapsulation device SA:V on syngeneic islet function in a rat omentum transplant model (**Figure 1B**).

**Figure 1.**
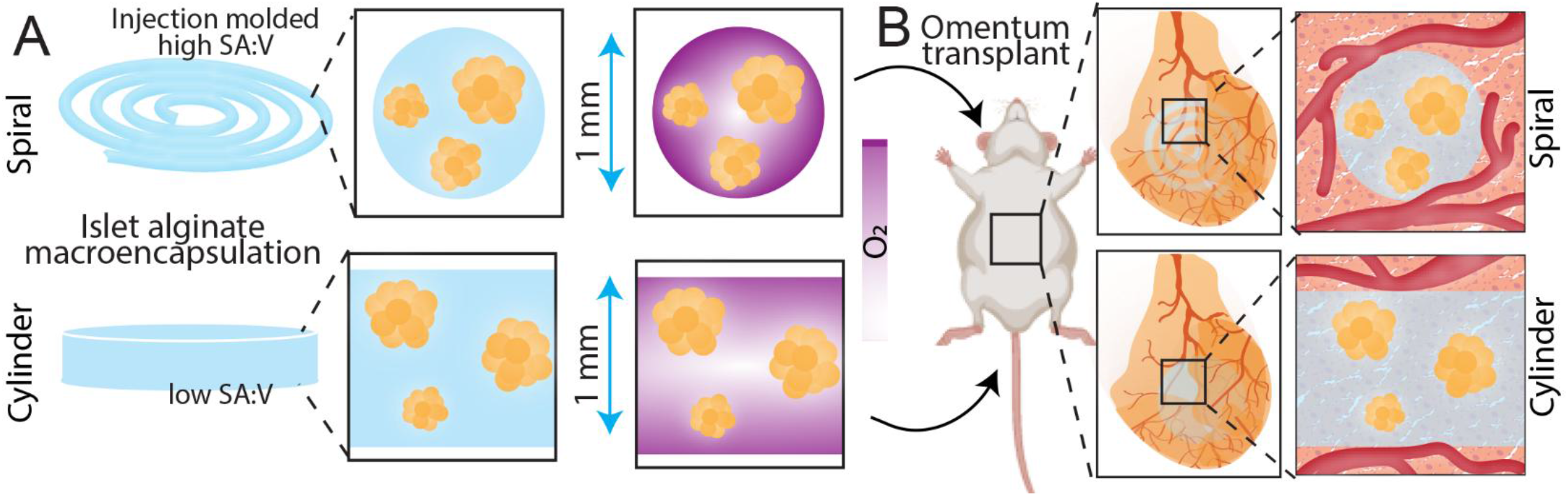
High SA:V macroencapsulation device design and transplantation strategy. (**A**) A hydrogel injection molded 1.5% (w/v) alginate spiral (high SA:V) geometry is compared against a traditional open-molded cylindrical (low SA:V) geometry of the same height (1 mm). (**B**) Syngeneic islet-containing macroencapsulation devices are delivered to the highly vascularized rat omentum site in diabetic recipients using a degradable, vasculogenic PEG hydrogel to promote vascularization at the device surface.

## 2. Materials and Methods

### 2.1. Materials

Chemicals were obtained from Sigma-Aldrich (St. Louis, MO), and cell culture materials were obtained from Thermo Fisher (Carlsbad, CA) unless otherwise noted. Peptides were obtained from GenScript (Piscataway, NJ) unless otherwise noted.

### 2.2. Animals

Male and female Lewis rats were purchased from Charles River Laboratory at 3 months old. All animal procedures were approved by the Arizona State University Institutional Animal Care and Use Committee and complied with relevant ethical regulations.

### 2.3. Macroencapsulation device fabrication and characterization

#### 2.3.1. Slow-gelling 1.5% alginate hydrogel fabrication

Slow-gelling alginate was fabricated using 3% alginate (UP-MVG, Novomatrix), 30 mM calcium carbonate (CaCO_3_), and gluconic δ-lactone acid (GDL), where varied concentrations of 40 mM, 60mM, 80 mM, and 100 mM GDL were used to determine the optimal crosslinking rate. Alginate and GDL solutions were made in DPBS -/- and stored at 4°C for 24 hours to allow for full solubilization. After 24 hours, CaCO_3_ was added to the alginate solution and mixed well. The GDL solution was added in a 1:1 ratio resulting in a final alginate concentration of 1.5% w/v and a final CaCO_3_ concentration of 15 mM. For photography purposes, various colors of food dye (U.S. Art Supply, San Diego, CA) were added to the GDL solution before gelation.

#### 2.3.2. Evaluation of hydrogel properties via rheometry measurements

Time studies were carried out to assess the gelation point via storage and loss moduli crossover for hydrogels. The hydrogel solutions were mixed and immediately injected onto a Physica MCR 101 rheometer (Anton Paar, Austria) at 25°C. Hydrogel storage and loss modulus were recorded over time (0.1% strain, 1 Hz) to display the crossover gelation point and complex viscosity of the sample before and after gelation. All GDL concentrations (40 mM, 60 mM, 80 mM, and 100 mM) were evaluated after being mixed with 3% alginate and 30 mM CaCO_3_ in a 1:1 ratio.

### 2.4. Computational modeling of islet loading density within macroencapsulation devices

A finite element modeling (FEM) approach was implemented on COMSOL Multiphysics. Diffusion was assumed to be governed by the diffusion equation

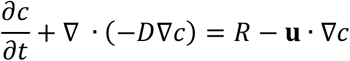

where c denotes the concentration (mol·m^-3^) and D the diffusion coefficient of oxygen through the hydrogel material (alginate), R the oxygen consumption rate of the islets (mol·m^-3^·s^-1^), **u** the velocity field (m·s^-1^), and ∇the standard nabla operator. For oxygen consumption, a Michaelis-Menten type consumption rate (R<0) was assumed.

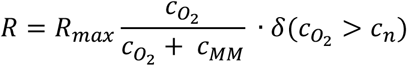

*R*_*max*_ is the maximum oxygen consumption rate, *c*_*MM*_ is the Michaelis-Menten constant, *c*_*n*_ is the oxygen concentration at which necrosis is assumed, and *δ* is a step down function to account for the cells that cease to consume oxygen once necrosis occurs. For these calculations, the oxygen concentrations of 20% and 7.23% were explored with a surface concentration value of 0.2 mol·m^-3^ and 0.0723 mol·m^-3^ respectively. These values represent relevant physiological oxygen concentrations associated with the omentum transplant site^29^. The alginate hydrogel material was assumed to have a diffusion coefficient of 2.5 ×10^−9^ m^2^·s^-1^ with respect to oxygen concentration. Various cell loading densities each corresponding to a density of 5 islet equivalents (IEQ)/μL, 10 IEQ/μL, 25 IEQ/μL, and 50 IEQ/μL were examined in the model to understand the relationship between cell loading and hypoxia-induced cell death. To calculate the area of islets in one cross section of the macroencapsulation device, the percentage of islet volume within one microliter of hydrogel was calculated and used to determine the percent islet area in one cross section. The average size of an islet was assumed to have a diameter of 150 micrometers. The islets were then evenly distributed thoughout the model with varying diameters representative of the typical islet size range. For each islet loading density group, six independent models were used to calculate the relationship between islet diameter and oxygenation. A maximum oxygen consumption rate of 0.034 mol·s^-1^·m^-3^ (per unit volume) was assumed. A Michealis-Menten constant of 1×10^−3^ mol·m^-3^ was assumed. For the step down function, COMSOL’s smoothed Heaviside function was used with the oxygen consumption rate reaching 0 at a concentration of 1×10^−4^ mol·m^-3^.

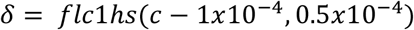

This model was solved as a time-dependent problem up to 3 days to correspond to *in vitro* assessments at 72 hours. All data displayed are results from day 3 of the study.

**Table 1.**
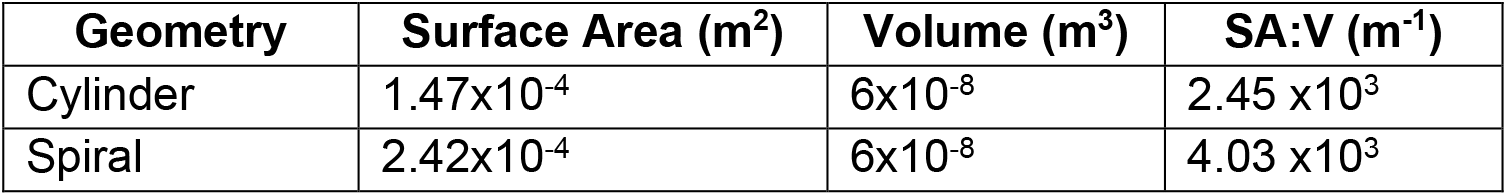
Summary of macroencapsulation device geometric qualities.

### 2.5. *In vitro* validation of computational models using pseudoislets

#### 2.5.1. Pseudoislet fabrication

Pseudoislets were fabricated as a cost-effective alternative to rat islets to assess the viability and function of various islet loadings within the spiral and cylindrical geometry alginate hydrogels. Non-treated petri dishes of 100 mm diameter were coated with 5 mL of anti-adherence rinsing solution (STEMCELL Technologies, Vancouver, CA) for 5 minutes at 37°C to prevent cell adhesion. The beta cell line INS1E, kindly gifted by Andres J. Garcia, was cultured in RPMI supplemented with 10% FBS, 1% penicillin-streptomycin, 1% HEPES buffer solution, 1% sodium pyruvate, and 0.09% β-mercaptoehtanol. INS1E cells were passaged and counted. The anti-adherence solution was aspirated from the petri dish, and the petri dish was rinsed with 5 mL of RPMI culture media and aspirated. Five million INS1E cells were added to the petri dish along with a final media volume of 12 mL. The cells were cultured for 4 days to allow for pseudoislet diameters comparable to rat islet diameters.

#### 2.5.2. Assessment of pseudoislet viability and function within macroencapsulation devices

Alginate (1.5% w/v) hydrogels were fabricated as described above with 30 mM CaCO_3_ and 100 mM GDL. Pseudoislets were collected, counted, and aliquoted to ensure a pseudoislet density of 5, 10, 25, and 50 islet equivalents (IEQ)/μL for each hydrogel cylinder and spiral. The cells were suspended in hydrogel solutions, injected into the injection mold or open cylinder mold, and allowed to gel for 5 minutes. The hydrogel constructs were carefully removed from their molds and washed with a 1.5% barium chloride (BaCl_2_) solution. The hydrogel constructs were cultured in well plates under standard conditions (5% CO_2_, 20% O_2_) as well as hypoxic conditions (5% CO_2_, 5% O_2_). At 72 hours post-construct fabrication, glucose stimulated insulin response (GSIR) was performed in static incubation and standard culture conditions in KREBS Buffer (15 mM NaCl, 4.7 mM KCl, 1.2 mM MgSO_4_, 2.5 mM CaCl_2_, 26 mM NaHCO_3_, and 0.2% BSA) with low (3 mM) and high (16 mM) glucose concentrations. Encapsulated pseudoislets were exposed to low glucose buffer for a 1 hour pre-incubation and 2 hour sequential low-high-low incubations. Samples were frozen at -80° C for later analysis via rat insulin ELISA (Mercodia, Uppsala, Sweden). AlamarBlue metabolic assay was mixed in a 1:10 ratio with the complete RPMI media per the manufacturer’s instructions and cultured with hydrogels for 4 hours prior to fluorescence measurement (ex/em 560/590 nm). Live/dead (calcein AM/ethidium homodimer) staining was performed per manufacturer’s instructions and imaged via a Leica SP8 White Light Laser Confocal microscope, housed in the Regenerative Medicine and Bioimaging Facility at Arizona State University.

### 2.6 Lewis rat islet Isolation and culture

Islets were isolated from male Lewis rats (Charles Rivers Lab) under protocols approved by the Institutional Animal Care and Use Committee (IACUC) at Arizona State University. Islets were obtained by mechanically and enzymatically digesting donor pancreata using DNase (Cat. No. 10104159001; Roche Diagnostics), Collagenase (Cat. No. 05989132001; Roche Diagnostics), and Thermolysin (Cat. No. 05989132001; Roche Diagnostics) as previously described^35^. The donor islets were then separated using density gradients (Cat. No. 99-692-CIS, Cat. No. 99-691-CIS, Cat. No. 99-815-CIS; Corning). Isolated rat islets were cultured overnight in CMRL 1066 media (Cat. No. 99-663-CV; Corning) with 10% fetal bovine serum, 1% penicillin-streptomycin, 1% L-glutamine, and 25 mM HEPES under standard incubation conditions (5% CO_2_ and 37°C) before transplantation. Islets were manually counted and quantified into IEQ. Islet purity was quantified by ditizone (DTZ) staining on the day of transplant. Islets stained with DTZ were considered functioning islets and non-functioning islets were considered part of the exocrine tissue. Rat islet purity was analyzed in FIJI using the following equation:

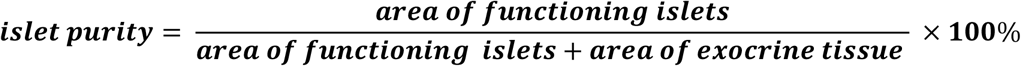

### 2.7. *In vitro* characterization of lewis rat islets

Alginate (1.5% w/v) hydrogels were fabricated as described above in 1 mm diameter spiral injection molds, 1 mm height cylindrical control hydrogels, or cultured in standard 2D conditions. Rat islets were isolated from male Lewis rats as described above. The islets were encapsulated 24 hours after isolation at a density of 10 IEQ/μL and cultured for 72 hours prior to assessment. GSIR, alamarBlue metabolic assay, and live/dead imaging were performed as described above.

### 2.8 Syngeneic lewis rat islet transplantation at omentum transplant site

Diabetes was induced in female Lewis rats by one injection of streptozotocin (STZ; Sigma Aldrich) at a dose ranging from 50 to 60 mg/kg. After three consecutive blood glucose readings greater than 350 mg/dL (3-6 days post STZ injection), the animals were considered diabetic and ready for transplantation. On the day of transplantation, islets were encapsulated in 1 mm diameter spiral alginate hydrogels or 1 mm height cylindrical alginate hydrogelsgroup of Unencapsulated islets served as a positive control. Islets were encapsulated at a density of 10 IEQ/μL resulting in a total of 600 islets per hydrogel construct. For the 1,200 IEQ dosage (6,000 IEQ/kg) this resulted in 2 spiral hydrogel constructs, 2 cylinder hydrogel constructs, or 1,200 unencapsulated IEQ into the omentum. For the 2,400 IEQ dosage (12,000 IEQ/kg)this dosage resulted in 4 spiral hydrogel constructs, 4 cylinder hydrogel constructs, or 2,400 unencapsulated IEQ per recipient. Animal anesthesia was induced with 5% isoflurane and maintained at 2% isoflurane. Primary rat islets were transplanted in the omentum. All devices were delivered with a thin layer of a vasculogenic, degradable hydrogel system established in previous studies^32^ to maximizwe vascularization and subsequently tissue oxygenation at the device surface. After surgery, blood glucose, body weight, and the general health conditions were monitored daily. Normoglycemia was defined as 250 mg/dL.

### 2.9 Intraperitoneal glucose tolerance test (IPGTT)

Animals were fasted overnight before the procedure as approved by IACUC at ASU. After blood glucose and body weight measurements, animals were injected with a 25% glucose-saline solution at a dose of 2 g/kg. Blood glucose was measured up to 240 minutes post-injection.

### 2.10 Histological assessment of explants

Retrieved grafts were fixed in 10% neutral-buffered formalin solution for 48 hours, transferred into 1x PBS, histologically processed with ethanol and embedded in paraffin. Resulting samples were sliced at 10 µm by microtome and stained using hematoxylin & eosin stain, Masson’s Trichrome, or an immunohistochemistry (IHC) panel. For IHC staining, insulin (rabbit anti-insulin, Cat. No. 701265 Thermo Scientific, 1:1000) was used followed by Alexa Flour 488 goat anti-rabbit and DAPI (Cat. No. 62248, Thermo Scientific, 1:1000 dilution). Fluorescent images were taken using a Leica SP8 confocal microscope. The thickness of the fibrotic response seen in H&E slides and the thickness of the collagen deposition seen in Masson’s Trichrome was analyzed using using image analysis and manual size annotation on an ECHO Revolve microscope. To measure vascularization, the distance of blood vessels from the surface of the implant in encapsulated groups and from the surface of the islet in unencapsulated groups was measured by manual annotation on an ECHO Revolve microscope.

### 2.11 Statistical analysis

In all studies, the results were expressed as mean ± SEM unless otherwise specified. Gelation time of 1.5% alginate hydrogel, macroencapsulation device average oxygen concentration results, and blood glucose results were analyzed by ordinary one-way ANOVA. pH of 1.5% alginate hydrogel, all rat islet GSIR index value data, all alamarBlue data, IPGTT results, quantification of the fibrotic response, quantification of vascularization, and pseudoislet and islet diameters were analyzed by one-way ANOVA with Kruskal-Wallis multiple comparison test. All pseudoislet GSIR index value data and pseudoislet alamarBlue data was analyzed by two-way ANOVA. All statistical analysis was conducted in GraphPad Prism 9.1 (GraphPad Software; San Diego, CA, US).

## 3 Results and Discussion

### 3.1 Computational modeling predicts islets in high SA:V geometry macroencapsulation devices have better viability and function

We hypothesized that islet macroencapsulation devices with a high SA:V ratio would significantly improve oxygen availability to encapsulated cells, enhance islet function and viability, and potentially increase the viable islet density loading limit within individual devices. To test this hypothesis, we used finite element modeling to evaluate various islet density loadings within two macroencapsulation devices of equal volume with either a high SA:V ratio (spiral geometry, 4.03 × 10^3^ m^-1^) or low SA:V ratio (cylinder geometry, 2.45 × 10^3^ m^-1^) (**Figure 2A**). A COMSOL model was developed to predict the local oxygen availability of individual encapsulated islets within alginate macroencapsulation devices at low (5 IEQ/µL), intermediate (10 IEQ/µL), and high (25 and 50 IEQ/µL) islet densities under standard culture oxygen conditions (20% O_2_, **Figure 2B-D**) and predicted *in vivo* oxygen conditions (7.23% O_2_, **Figure 2E-G**). In the 20% oxygen boundary condition model, reflective of standard *in vitro* culture conditions, 5 and 10 IEQ/µL loading densities resulted in significantly higher islet oxygen concentration than 25 or 50 IEQ/µL for both spiral and cylinder geometries (**Figure 2C**), and islet oxygen availability decreased with increasing islet diameter (**Figure 2D**). Further, islet loading densities of 25 and 50 IEQ/µL in the cylindrical device demonstrated average oxygen levels significantly lower than the corresponding densities in the spiral geometry (**Figure 2C**), and lower than 0.03 mol/m^3^ (**Figure 2D**), which is the oxygen level below which insulin secretion is impaired.

**Figure 2.**
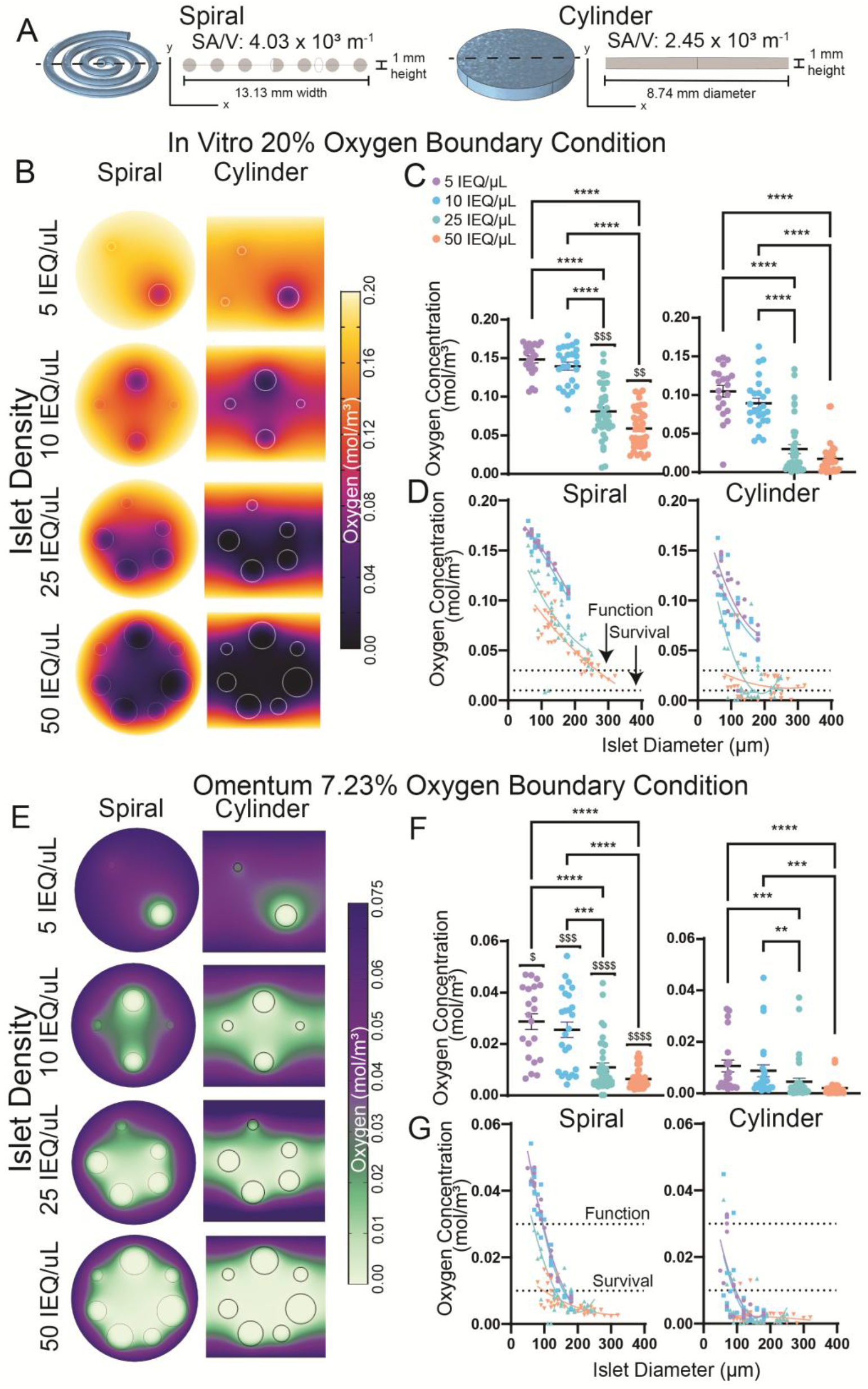
Computational modeling demonstrates macroencapsulation devices with greater SA:V have improved islet survival and function at physiological oxygen concentrations, allowing higher islet density loading. (**A**) Macroencapsulation device dimensions and cross-section. Oxygen boundary conditions were evaluated for standard in vitro culture (20% O_2_, B-D), and estimated vascularized omentum transplant site (7.23% O_2_, E-G) at a range of islet density loadings. Oxygen distributions are demonstrated via (B,E) cross section of spiral arm (1 mm diameter) and cylinder x-y plane (1 mm height), (C,F) plotting individual islet oxygen concentration, and (D,G) distribution of islet oxygen concentration by islet diameter. Dashed lines indicate thresholds for islet function and survival. Scale bars = 1 mm. Statistical analysis by ordinary one-way ANOVA, $ cell density compared within oxygen environment spiral vs cylinder. One symbol P < 0.05, two symbols P < 0.01, three symbols P < 0.001, four symbols P < 0.0001. Error bars = SEM.

We next modeled macroencapsulation devices in a 7.23% oxygen environment (**Figure 2E-G**), which is equivalent to physiological oxygen within the vascularized omentum transplant site, a level slightly higher than the subcutaneous transplant site (5.92% oxygen)^32,36^. This boundary condition exhibited similar trends to the 20% oxygen model, with 5 and 10 IEQ/µL loading densities exhibiting significantly higher islet oxygen concentration than 25 or 50 IEQ/µL for both spiral and cylinder geometries (**Figure 2F**). In addition, average values for all islet loadings in the cylindrical device of the 7.23% oxygen model drop below the 0.01 mol/m^3^ threshold (dashed line, **Figure 2F**) for islet survival while the average values for the spiral device at 5 and 10 IEQ/µL loadings remain at the 0.03 mol/m^3^ threshold (**Figure 2G**). Further, the spiral geometry demonstrated significantly higher average oxygen concentration at all islet loading densities relative to the cylindrical geometry (**Figure 2F**).

In sum, computational oxygen modeling demonstrates that the high SA:V spiral device preserves islet viability and function at higher islet loadings in *in vitro* oxygen conditions and at low islet loadings (5-10 IEQ/μL) under physiological oxygen concentrations, whereas traditional low SA:V device designs exhibit significantly reduced oxygen availability. Further, large diameter islets have higher oxygen needs and are particularly prone to necrotic cores^37^, and islets with diameters greater than 200 µm exhibited poor survival regardless of islet density or device geometry. In all models, increasing islet diameter negatively correlated with oxygen concentration. Overall, 5 IEQ/μL and 10 IEQ/μL spiral islet loadings behaved comparably at both 20% and 7.23% oxygen levels, suggesting up to 10 IEQ/μL is a suitable spiral macroencapsulation device loading to preserve device function.

### 3.2 Slow-gelling alginate pH and mechanical property optimization for hydrogel injection molding complex macroencapsulation device geometries

After computational oxygen modeling demonstrated that high SA:V geometries such as the spiral promote greater islet survival and function, we next sought to fabricate these devices to validate our models. We recently developed a hydrogel injection molding method to fabricate complex hydrogel macroencapsulation geometries such as the high SA:V spiral using a variety of natural^29^ and synthetic^30^ materials (**Figure 3**). To fabricate alginate spirals via injection molding, we adapted a slow-gelling alginate crosslinking method^38^ using CaCO_3_ (30 mM) and GDL (100 mM), which allows time for alginate injection into the mold prior to gelation (**Figure 3A**). In this reaction, Ca^2+^ ions are released from CaCO_3_ through intereaction with free protons released by the hydrolysis of GDL. This process results in sufficient time to gelation to allow injection into the mold before crosslinking (**Figure 3B**), and the alginate crosslinking is reinforced after extraction from the mold with a barium ion crosslinking bath to ensure greater hydrogel stability.

**Figure 3.**
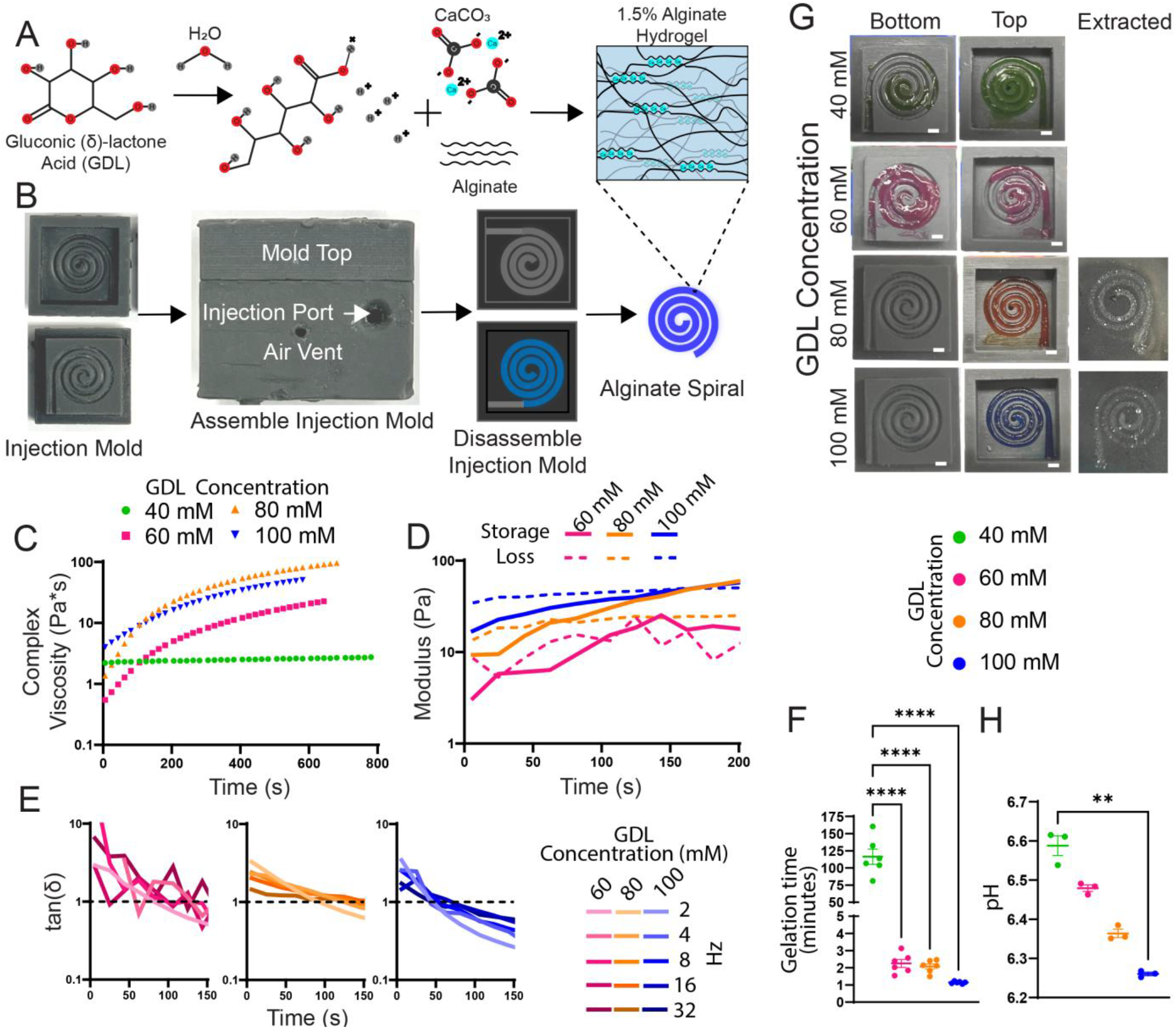
Slow-gelling alginate composition optimization for hydrogel injection molding suitability via crosslinking time, mechanical stiffness, and pH. (**A**) Gluconic (δ)-lactone (GDL) hydrolyzes gradually to gluconic acid in solution, releasing protons to liberate calcium ions from calcium carbonate, which can then crosslink alginate. (**B**) Slow gelation of alginate enables injection into molds to generate complex geometries. (**C**) Complex viscosity (Pa*s), (**D**) storage and loss modulus, and (**E**) tan(δ) were characterized by rheometry to determine hydrogel suitability for injection molding. (**F**) The gelation time of each GDL concentration was validated manually, and (**G**) hydrogel formation in molds and extraction were characterized. (**H**) GDL concentration influence on alginate pH was characterized. Scale bars = 1 mm. Error bars = SEM. Gelation time was analyzed by ordinary one-way ANOVA. pH analyzed by one-way ANOVA with Kruskal-Wallis multiple comparison test. *P < 0.05, ****P<0.0001.

We previously demonstrated that human islets encapsulated in this alginate formulation function comparably to unmanipulated islets ^29^; however, high GDL concentrations reduce alginate pH, and we aimed to minimize pH reduction in the gel to limit encapsulated cell stress. As such, to ascertain whether GDL could be reduced, we characterized the gelation point and mechanical properties of our slow-gelling alginate using rheometry, evaluating a range of GDL concentrations (40, 60, 80, and 100 mM). 1.5% (w/v) alginate gels were formulated with 30 mM CaCO_3_, which our previous work demonstrated was the concentration required to ensure maximal crosslinking of a 1.5% alginate hydrogel^29^. First, we characterized temporal hydrogel complex viscosity after mixing alginate, GDL, and CaCO_3_ components. We previously demonstrated that limiting hydrogel complex viscosity below 100 Pa*s before and during injection avoids cytotoxic pressure levels within the injection mold. All formulations with the GDL concentrations exhibited complex viscosities below 100 Pa*s for longer than 10 minutes (**Figure 3C**) suggesting all formulations are suitable for injection molding within the timeframe required to inject cell/hydrogel solutions into the mold. 60-100 mM GDL solutions increased in complex viscosity over time, whereas 40 mM GDL exhibited no change in viscosity over time, consistent with no gelation within the 10-minute time frame.

We next evaluated the temporal storage and loss modulus of 60, 80, and 100 mM GDL slow gelling alginates (**Figure 3D**) to determine the temporal change in hydrogel stiffness and identify the storage and loss modulus crossover point, which indicates gelation point. The 40 mM GDL group was not included due to its gelation time exceeding our desired window to maintain cell viability (∼10-15 min). We found that the 80 mM and 100 mM GDL concentrations exhibited similar storage moduli of 53.5 and 53.7 Pa respectively around 3 minutes post-mixing, while 60 mM GDL concentration exhibited lower storage moduli of 19.2 Pa value at 3 minutes. The lower storage modulus plateau value for the 60 mM GDL group may be a result of an insufficient release of calcium from CaCO_3_ to generate stable alginate crosslinking.

Erratic storage and loss modulus measurements for the 60mM GDL group made it challenging to determine the accurate crossover point and estimate gelation time. As such, we aimed to more accurately assess the gelation point of each GDL group by (1) rheometry and (2) injectability testing. To evaluate gelation point via rheometry, we plotted the loss tangent (tan δ) of the 60 mM, 80 mM, and 100 mM GDL alginate formulations for multiple frequencies over time (**Figure 3E**). We found that alginate formulations with 60 mM, 80 mM, and 100 mM GDL concentrations exhibited gelation points of 132, 123, and 50 s, respectively. To test the limit of injectability, we measured alginate formulation gelation by determining the point at which hydrogels were no longer injectable, finding gelation points of 116 minutes, and 145, 127, and 74 seconds for 40, 60, 80, and 100 mM GDL, respectively (**Figure 3F)**. Gelation point measurement using rheometry and subjective injectability were remarkably well aligned for 60 mM GDL concentrations and above. We also evaluated the integrity and extractability of injection molded alginate hydrogels, and found with GDL concentrations of 40 mM and 60 mM formed incomplete spiral hydrogels within the injection mold, whereas GDL concentrations of 80 mM and 100 mM GDL formed full spiral geometries that were easy to extract from the molds (**Figure 3G)**.

Finally, we evaluated the pH of the various alginate formulations before gelation to determine if the increased concentrations of GDL resulted in a potentially cytotoxic pH (**Figure 3H**). All pH values fell within a pH of 6.2 and 6.7 which is less than the pH of 6.8 needed to maintain normal intracellular pH^39^; however, brief exposures to this lower pH during encapsulation do not lead to long-term effects ^29^. We demonstrated the 40 mM GDL concentration, which possessed the highest pH, was unsuitable for injection molding due to its stiffness and gelation time, and the remaining concentrations of GDL possessed pH values that were slightly lower. Thus, we determined that the pH values of the 60 mM, 80mM, and 100 mM hydrogel formulations posed a relatively equal viability risk to encapsulated islets. Cumulatively, the 100 mM GDL group demonstrated gelation times within the desired window of 1-5 minutes, an increased stiffness relative to other concentrations, and successful spiral fabrication within the injection mold, and we chose to proceed with this formulation for all future studies.

### 3.3 High SA:V ratio macroencapsulation device geometry improves islet viability and function *in vitro* relative to traditional device designs

We next sought to validate our computational models of environmental oxygen concentration and islet density impact on cell function and viability within macroencapsulation devices *in vitro* (**Figure 4** and **5)**. We first fabricated pseudoislets using the insulin responsive INS1E rat-derived β cell line, generating cell clusters with an average diameter 96.2 µm (**Supplementary Figure 1**), to evaluate a range of islet loading densities from 5-50 IEQ/µL under 20% and 5% oxygen conditions (**Figure 4A**). Pseudoislets were chosen as a cost-effective islet substitution due to their ease of fabrication and mass production, and their capacity to mimic islet hypoxic cores. We evaluated function and viability at 72 hours to allow suitable time for the development of hypoxic cores.

**Figure 4.**
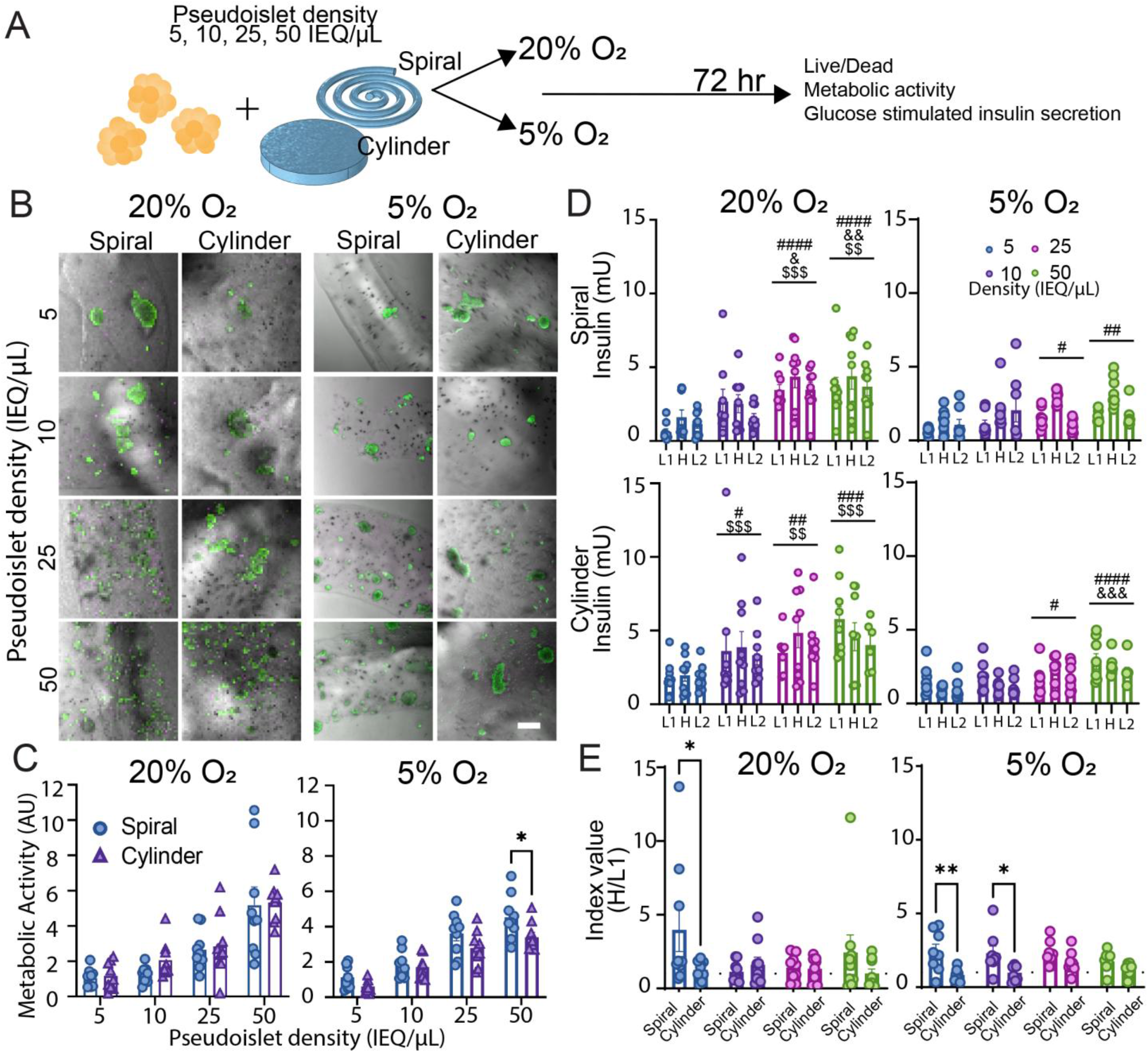
Encapsulated pseudoislets cultured at physiological oxygen levels validates computational models of islet viability and function in vivo. (A) Schematic illustrating in vitro macroencapsualted pseudoislet conditions and assessment. (B) Live (green)/dead (magenta) confocal imaging, (C) alamar blue metabolic activity, and (D) glucose stimulated insulin response assay of encapsulated pseudoislets after 72 hours of culture in 20% and 5% oxygen condiitons. (E) GSIR index values, the ratio of high glucose to low glucose insulin secretion. Live/dead n = 3 gels/group, n = 9 gels/group (**C-E**). Replicates were pooled from three independent experiments. Error bars = SEM. Scale bars = 200 µm. GSIR, index values, and alamar blue data analyzed by two-way ANOVA: # vs 5 IEQ/µL, & vs 10 IEQ/µL, $ vs 5% O_2_, same cell density. One symbol P < 0.05, two symbols P < 0.01, three symbols P < 0.001, four symbols P < 0.0001.

Live/dead staining of pseudoislets (**Figure 4B**) within hydrogel macroencapsulation devices indicate high viability of pseudoislets across all density distributions and oxygen conditions. Pseudoislet metabolic activity at 20% O_2_ (**Figure 4C**) demonstrated comparable values between spiral and cylindrical hydrogel geometries at all islet loading densities, but cylinder devices exhibited a trend of reduced metabolic activity at higher density loadings at 5% O_2_, with significantly reduced metabolic activity at the 50 IEQ/µL loading relative to the spiral geometry. We also evaluated pseudoislet function using the glucose stimulated insulin release (GSIR) assay, where low (3 mM), high (16 mM), and second low (3 mM) glucose incubations should produce a biphasic insulin response (**Figure 4D**). Total insulin secretion levels increased with increasing cell loading in a dose dependent manner in both geometries in 20% oxygen condition. Overall insulin secretion was reduced in 5% O_2_ conditions compared to 20% O_2_ at 25 and 50 IEQ/μL loadings in spiral devices and at 10-50 IEQ/μL loadings in cylindrical devices. The GSIR index value (high glucose value divided by 1^st^ low glucose value) is an additional measure of insulin responsiveness and islet health (**Figure 4E**). The spiral geometry with a 5 IEQ/µL loading density exhibited a significantly greater GSIR index value than the cylinder geometry at 20% O_2_, and significantly greater than the cylinder at 5% O_2_ for both 5 IEQ/µL and 10 IEQ/µL loading density. This aligns with our 5% O_2_ COMSOL modeling where the oxygen concentration at both 5 IEQ/µL and 10 IEQ/µL loading densities were above the 0.03 mol/m^3^ oxygen threshold in the spiral device but below this critical threshold in the cylindrical device. Based on pseudoislet viability and function data at 5% and 20% O_2_, we selected the 10 IEQ/µL loading density for all future experiments.

We next validated our pseudoislet observations with primary rat islets. We examined the viability, metabolic activity, and function of primary rat islets in our 1.5% alginate spiral and cylinder macroencapsulation devices at the 10 IEQ/μL loading and 20% O_2_. A high degree of islet viability was observed for both macroencapsulation geometries (**Figure 5A**), with the cylinder group exhibiting fewer healthy-looking islets in the central region of the device. The metabolic activity of the cylindrical macroencapsulation device was significantly lower than the spiral device, which demonstrated comparable metabolic activity to unencapsulated islets (**Figure 5B**). Islet insulin responsiveness via GSIR assay demonstrated a biphasic insulin response in spiral and unencapsulated groups, but not the cylinder macroencapsulated group (**Figure 5C**). While GSIR index values for cylinder (0.996) trended lower than spiral (1.363) and unencapsulated (1.444), this difference was not statistically significant (P = 0.1044) (**Figure 5D**).

**Figure 5.**
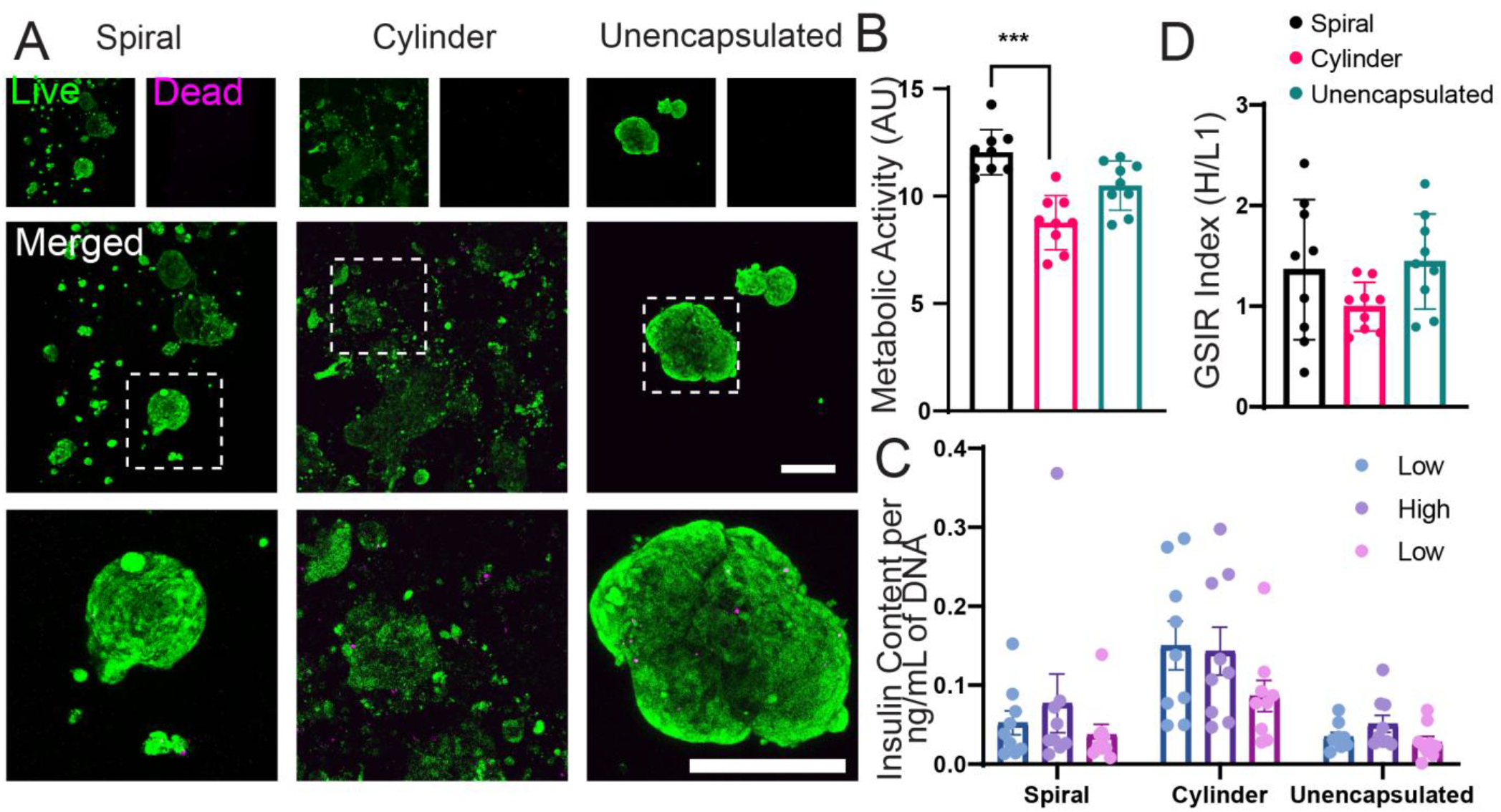
High SA:V geometry macroencapsulated rat islets perform comparably to unmodified islet in vitro. (A) Live (green)/dead (magenta) confocal imaging of rat islets encapsulated in spiral geometry and cylindrical geometry compared against unencapsulated rat islets under standard culture conditions. (B) Metabolic activity of the encapsulated and unencapsulated rat islets was evaluated via alamarBlue assay, and insulin responsiveness (C) and index values (D) evaluated via GSIR. n = 9 gels/group for alamarBlue and GSIR assays. n = 3 gels/group for live/dead imaging. Replicates were pooled from three independent experiments. Metabolic activity and GSIR index values analyzed by one-way ANOVA with Kruskal-Wallis multiple comparison test. *P < 0.05, ***P < 0.001. Error bars = SEM. Scale bars = 200 µm.

Three separate preparations of rat islets were pooled from three independent islet isolations which may result in variations among individual data points (**Supplementary Figure 2**). However, all preparations of rat islets exhibited comparable results and insulin responsiveness (**Supplementary Figure 2F**). We also examined islet purity on the effects of islet function and found no significant difference between the islet purity of the individual batches (**Supplementary Figure 2B**). The individual batches 1, 2, and 3 possessed an average islet purity in the range of 42% to 48%, which is representative of the mean islet purity of 45% observed in human islet transplantation^40^. The higher metabolic activity observed in the spiral group relative to the cylinder group is likely due to shorter diffusion distances, resulting in improved oxygen and nutrient transport (**Supplementary Figure 2D**). Overall, our *in vitro* data matches *in silico* predictions that a detriment to islet function and viability becomes evident at higher encapsulation densities and lower oxygen concentrations, and that densities up to 10 IEQ/µL can support functional encapsulated islets within high SA:V device geometries.

### 3.4 High SA:V spiral macroencapsulation device geometry demonstrates improved syngeneic islet graft function in a diabetic rat omentum transplant model relative to traditional low SA:V design

*In silico* modeling and *in vitro* observations indicated that the spiral geometry preserved greater viability and function of encapsulated cells, and that the 10 IEQ/µL density was the highest density to preserve islet viability and function. We next sought to validate these observations *in vivo* in a diabetic rat model and evaluate the impact of macroencapsulation device geometry on syngeneic islet function in an omentum transplant site (**Figure 6-8, Supplementary Figure 3-6**). We chose the rat omentum model due to its comparable anatomical structure to the human omentum, and a foreign body response that is more comparable to human than the mouse model^41^. Lewis rat islets were encapsulated in spiral or cylindrical alginate hydrogels at 600 IEQ per device resulting in an islet loading density of 10 IEQ/µL; control recipients received unencapsulated islets at the same dosage. Devices or unencapsulated islets were transplanted in the rat omentum with a degradable vasculogenic hydrogel that previously demonstrated the capacity to maximize vasculature at encapsulation device surface^32–34^ (**Figure 1**). This vasculogenic hydrogel degrades within 2-4 weeks and demonstrated improved unmodified and encapsulated islet engraftment in the omentum-equivalent murine epididymal fat pad site ^32–34^.

**Figure 6.**
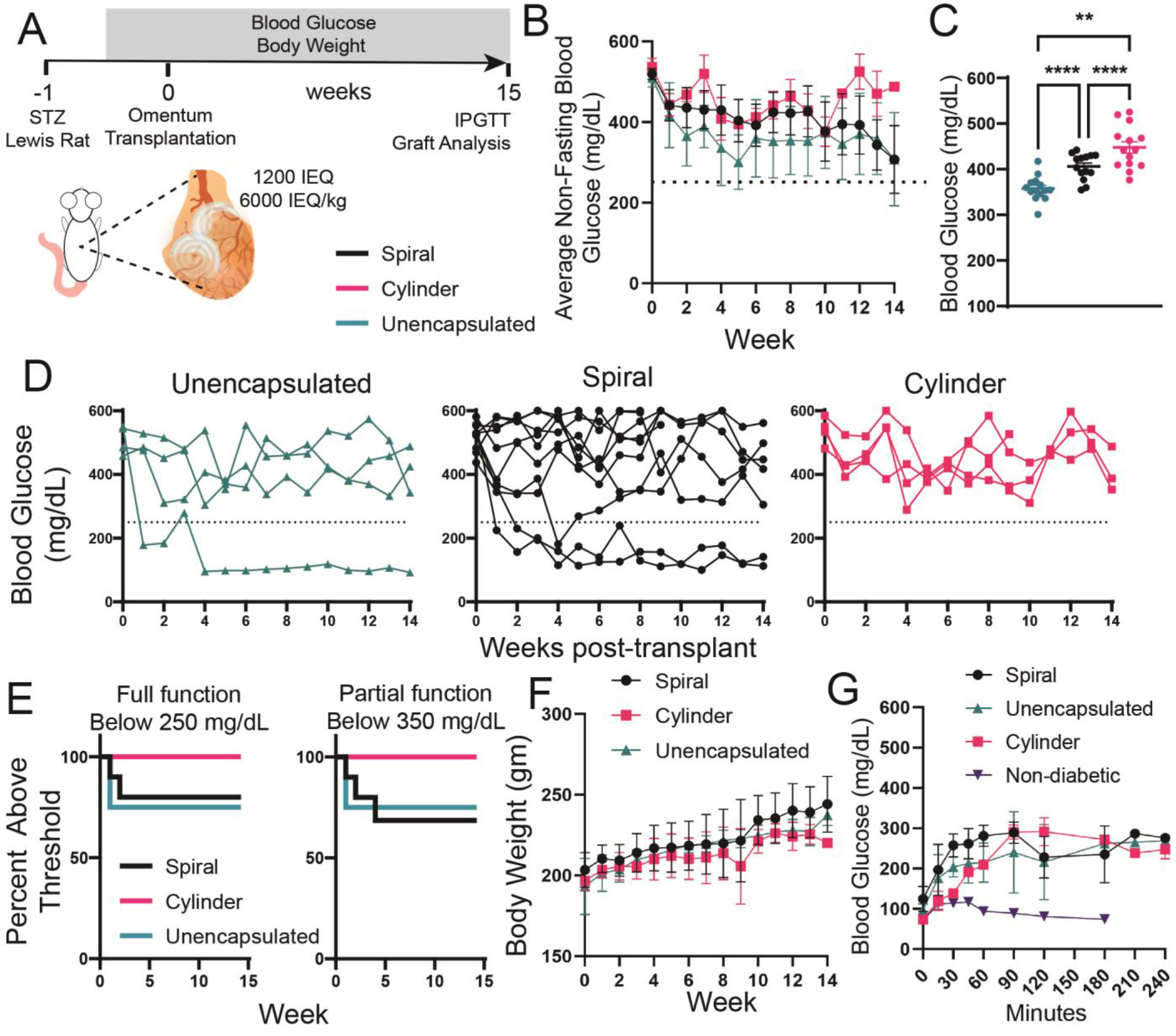
Subcurative spiral encapsulated islet dosage (6,000 IEQ/kg) results in comparable diabetes reversal rate to unencapsulated islets in rat omentum transplant model. (**A**) Schematic illustrating the experimental timeline. (**B**) Weekly average of nonfasting blood glucose level measurements, (**C**) pooled nonfasting blood glucose averages after week 1, and (**D**) individual rat blood glucose levels. (**E**) Survival curves for diabetes reversal (250 mg/dL) and partial graft function (350 mg/dL). (**F**) Average body weight measurements post-transplant. (**G**) Intraperitoneal glucose tolerance test (IPGTT) results during week 14. n = <u>4-8</u> for each transplant group. For IPGTT, n = 4 for spiral device, n = 2 for cylinder group, n = 2 for unencapsulated group, and n = 3 for non-diabetic group. Blood glucose results were analyzed by ordinary one-way ANOVA. **P < 0.01, ****P < 0.0001. Error bars = SEM.

We first evaluated a subcurative dosage of 1200 IEQ (6,000 IEQ/kg) (**Figure 6A**), which rarely reverses diabetes in syngeneic rat or allogeneic human islet transplantation ^26,42^. We chose this subcurative dosage as it would be comparable to a typical human islet transplantation from a single donor. Further, our islet preparation purity averaged 37.62% (**Supplementary Figure 4**), which is comparable to typical islet purities achieved for human islet transplantation preparations^43^. Previous studies have demonstrated that lower islet preparation purities increase the number of islets required to obtain function *in vivo*^26^. Average nonfasting blood glucose values in this cohort remained above the normoglycemia cutoff of 250 mg/dL for all groups (dashed line, **Figure 6B**), and we found that average blood glucose values after week one for cylinder recipients (450.6 mg/dL) were significantly greater than spiral (401.6 mg/dL) and unencapsulated (356.1 mg/dL), and spiral was significantly greater than unencapsulated (**Figure 6C**). No cylinder macroencapsulation device recipients achieved normoglycemia, whereas 25% of spiral and unencapsulated recipients achieved normoglycemia within 3 weeks (**Figure 6 D, E**). Additionally, 25% and 30% of unencapsulated and spiral recipients achieved average blood glucose levels below 350 mg/dL within 5 weeks of transplant, respectively, indicating at least partial function of grafts, whereas cylinder devices did not exhibit any measurable partial function by this measure (**Figure 6E**). In human islet transplant studies, a 6,000 IEQ/kg dose is typically sufficient to provide only partial function, reducing insulin requirements by about half ^42^. Throughout the 15 weeks, body weights of all groups increased comparably (**Figure 6F**). Finally, intraperitoneal glucose tolerance tests (IPGTT) were conducted on select recipients on week 15 (**Figure 6G, Supplementary Figure 3)**, and all groups demonstrated poorer glucose responsiveness relative to nondiabetic controls. Interestingly, all groups recorded fasted blood glucose readings within the normoglycemic range, providing further evidence of partial graft function.

We next evaluated whether functional differences between spiral and cylinder encapsulation geometries were maintained at a higher marginal dosage of 12,000 IEQ/kg (2400 IEQ total, 4 spirals or cylinders containing 600 IEQ each) (**Figure 7A**). In human islet transplantation, this dose is sufficient for ∼50% diabetes reversal rate, and partial graft function in the remaining patients^42^. Our islet preparation purity for these transplants averaged 28.27% (**Supplementary Figure 5**) which is lower than the purity average of the 6,000 IEQ/kg dosage cohort. Average nonfasting blood glucose values remained above the normoglycemia cutoff of 250 mg/dL for the spiral and cylinder groups while the unencapsulated recipients achieved normoglycemia within 2 weeks (**Figure 7B**). Pooling average nonfasting blood glucose values for each group from week 1 to 14 demonstrates that unencapsulated islets had significantly lower average blood glucose levels (223.3 mg/dL) than both macroencapsulation device geometries, and spiral device blood glucose levels (347.3 mg/dL) were significantly lower than that of cylinder devices (397.2 mg/dL) (**Figure 7C**). No spiral or cylinder macroencapsulation devices achieved normoglycemia, whereas 50% of unencapsulated recipients achieved normoglycemia within 1 week (**Figure 7D, E)**. Additionally, 80% and 75% of unencapsulated and spiral recipients achieved average blood glucose levels below 350 mg/dL within 6 weeks of transplant, respectively, indicating at least partial function of grafts, compared to only 50% of cylinder recipients which was statistically different from the unencapsulated group (**Figure 7E**). Throughout the 15 weeks, body weights of all groups increased comparably (**Figure 7F**). Finally, IPGTTs were conducted on select recipients on week 15 to evaluate graft glucose responsiveness (**Figure 7G, Supplementary Figure 3**), with insulin responsiveness reflecting average blood glucose values (**Figure 7C**). All groups demonstrated limited insulin responsiveness, as average blood glucose measurements did not return to baseline after 2 hours. However, the unencapsulated group exhibited significantly better insulin responsiveness than the cylinder group. Across all groups, we observed a decrease in average blood glucose values after increasing the islet dosage from 6000 IEQ/kg **(Figure 6C**) to 12,000 IEQ/kg **(Figure 7C**), with encapsulated group averages decreasing 54.3 and 53.4 mg/dL for spiral and cylinder devices, respectively, and 132.8 mg/dL in the unencapsulated group.

**Figure 7.**
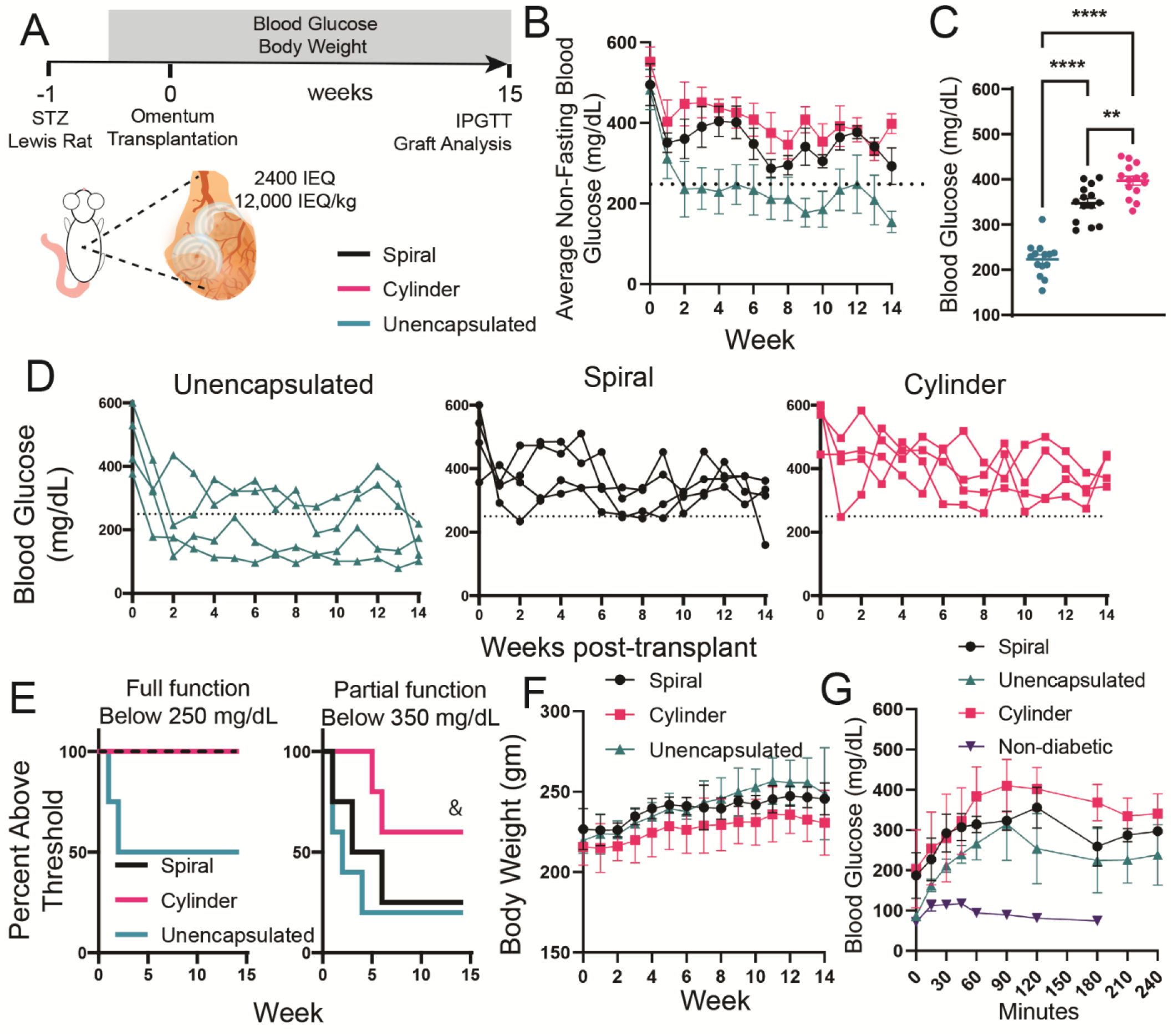
Marginal islet transplant dosage (12,000 IEQ/kg) demonstrates improved blood glucose control in high SA:V encapsulation device relative to low SA:V devices. (**A**) Schematic illustrating the experimental timeline. (**B**) Group averages and (**C**) overall averages after week 1 of non-fasting blood glucose level measurements performed 3 times per week. (**D**) Individual rat non-fasting blood glucose levels over 14 weeks. (**E**) Survival curves for diabetes reversal (250 mg/dL) and partial graft function (350 mg/dL). (**F**) Average body weight measurements post-transplant. (**G**) Intraperitoneal glucose tolerance test (IPGTT) results during week 14. Intraperitoneal glucose tolerance test (IPGTT) results during week 14 (**G**). n = 4 for each group. Non-diabetic group n = 3. Blood glucose results were analyzed by ordinary one-way ANOVA. **P < 0.01, ****P < 0.0001. &P < 0.05 vs unencapsulated group by Gehan-Breslow-Wilcoxon test. Error bars = SEM.

Finally, we evaluated local islet and tissue remodeling within macroencapsulated and control grafts via histological analysis (**Figure 8**). H&E staining (**Figure 8A**) of the explanted macroencapsulation devices shows detectable islets within devices and unencapsulated controls, as well as substantial tissue remodeling around alginate devices. Similarly, Masson’s Trichrome staining (**Figure 8B**) demonstrates noticeable collagen deposition (blue) around macroencapsulation devices, with limited deposition near unencapsulated control islets. Insulin positive islets were identifiable in all macroencapsulation devices and cell-only controls, with comparable morphology to nondiabetic pancreatic controls (**Figure 8C**). We next quantified the thickness of tissue on the surface of devices or in proximity to unencapsulated islets, and found both alginate geometries resulted in significantly greater tissue infiltration and remodeling at the device surface than in proximity to unenecapsulated islets, though the values for the two different geometries were comparable (**Figure 8D**). We next used collagen deposition in the graft, either at the device surface or around engrafted unencapsulated islets, as a measure of fibrosis (**Figure 8E**). We found only cylinder devices had a signficiant increase in collagen deposition relative to spiral and unencapsulated controls. Studies have shown that even the most biocompatible materials result in approximately 100 µm thick fibrotic tissue response after about 1 month^44,45^, which is similar to the response seen on our implanted devices. Finally, we quantified the distance of blood vessels in the graft from the device surface, and found that between 72-75% of blood vessels in the field of view were within 100 µm of cylinder or spiral device surface (**Figure 8F**). Finally, we performed H&E and insulin/DAPI immunostaining on explanted pancreata from graft recipients in **Figure 6** and **7** to confirm that recipient pancreata did not have residual function (**Supplementary Figure 6**). No islets were observed in the sectioned pancreata, confirming decreased blood glucose levels were caused by the implanted islets and not recovery of streptozotocin-treated islets.

**Figure 8.**
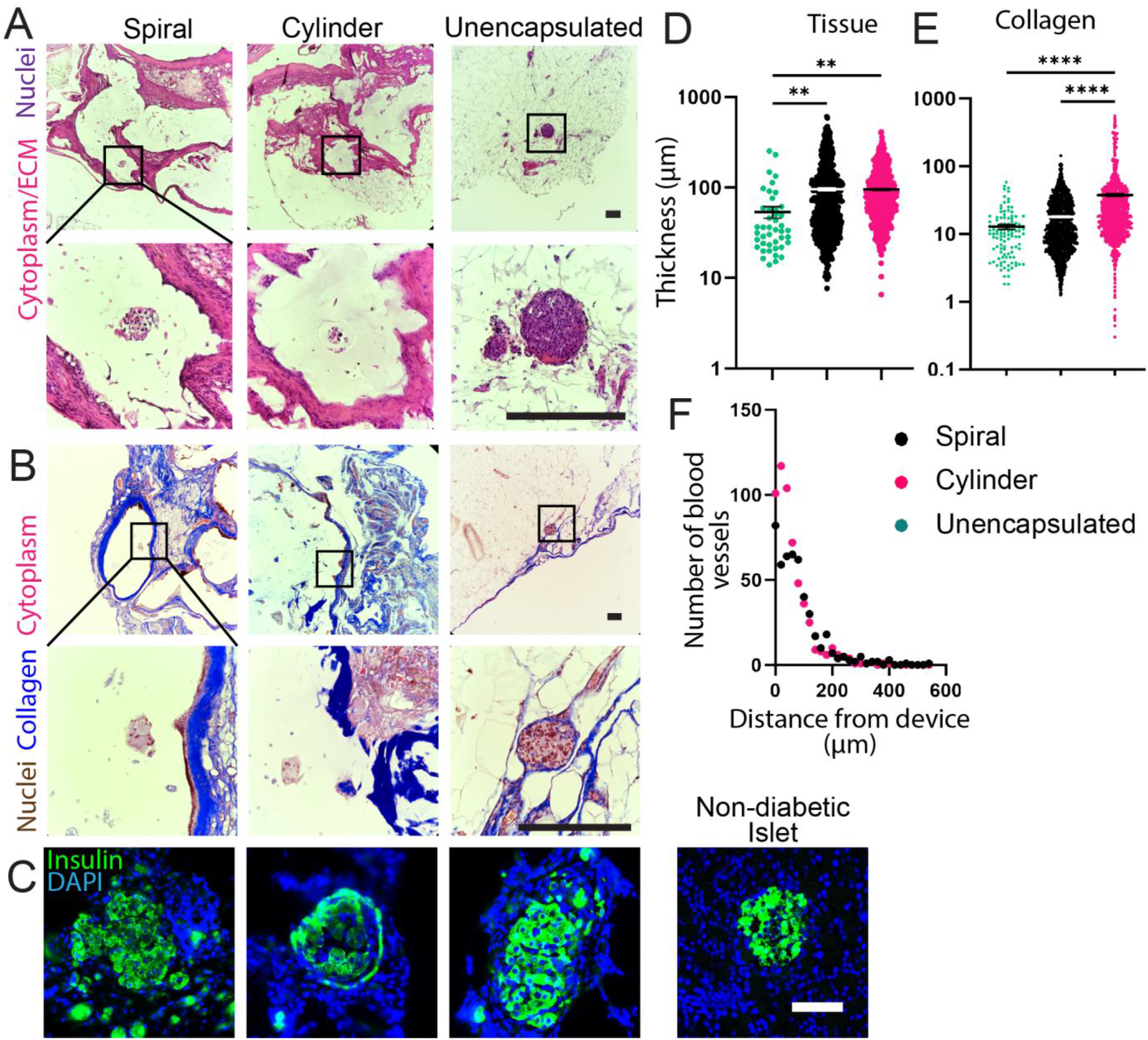
Macroencapsulated islet grafts are insulin-positive with the majority of blood vessels within 100 µm of the device surface. Macroencapsulation devices and unencapsulated control omentum grafts were retrieved on week 15 and stained for (**A**) H&E (cytoplasm/ECM in pink), (**B**) Masson’s trichrome (collagen in blue), and (**C**) insulin (green). Fibrotic response was quantified based on (**D**) tissue thickness in H&E sections and (**E**) collagen thickness in Masson’s Trichrome. Vascularization of devices and islets was examined by measuring the distance of blood vessels from the device and islet surface. Scale bar in H&E (**A**) and Masson’s Trichrome (**B**) is 200 µm. Scale bar in IHC (**C**) is 100 µm. Quantification of tissue and collagen thickness (data pooled from n = 3-6 biological replicates across both IEQ dosages) was analyzed by one-way ANOVA with Kruskal-Wallis multiple comparison test. *P < 0.05, **P < 0.01, *** P < 0.001, ****P < 0.0001. Error bars = SEM.

Overall, our unencapsulated omentum-transplanted islet reversal rates align with human islet transplantation reversal rates^42^, whereas spiral or cylinder encapsulated islets demonstrate some impairment, with spiral macroencapsulation devices demonstrating significantly improved blood glucose control relative to lower SA:V cylinder devices. The data suggest that macroencapsulated islet function can be improved through geometric design, though a clear limitation is that higher cell numbers may be required than no encapsulation to achieve normoglycemia. The data here align with previous studies delivering macroencapsulation devices to the omentum ^32,46^, and studies which demonstrated that encapsulation may require islet numbers greater than 12,000 IEQ/kg to reverse T1D^47^. Traditional microcapsules contain 2-3 IEQ/capsule; at a standard diameter of 1 mm per typical capsule, traditional microencapsulation results in an islet density of ∼5 IEQ/µL, which is half of the density we used in our macroencapsulation devices. Given an encapsulated islet reversal dosage of 15,000 IEQ/kg for a 70kg average man, this results in a minimal of 283 mL of encapsulated cell volume, including void space between capsules. This illustrates the limitations of microencapsulation transplant site, as delivering microcapsules to a site like the omentum or subcutaneous space would require a 38×38 cm area to deliver a double layer of capsules. By contrast, our clinical scale spiral designs of 2 mL each would result in 105 mL total volume and a footprint of 12.5×12.5 cm at the same dosage and 10 IEQ/µL density, which provides significantly greater flexibility in transplant site.

Additionally, while it was an advantage that our islet preparations had comparable purity rates to standard human islet preparations for transplantation, which results in a more accurate assessment of potential clinical outcomes, this inevitably led to a higher cell density within the devices than intended. For example, an islet purity of 30% and islet loading of 10 IEQ/µL would lead to three times as much tissue as the number of islets, which would inevitably impact oxygen consumption and transport, and likely contributed to poorer function in vivo than a purer preparation; however, the impact of this additional tissue on graft function is difficult to predict, as acinar tissue typically demonstrates 100-fold lower oxygen consumption rate than islets^48^.

The higher cell number requirement for encapsulated cell therapies is a significant clinical burden when sourcing cadaveric donor islets, as limited donor pancreata are available for islet transplantation^49^. However, the recent development of stem cell-derived insulin secreting cell sources, currently accelerating through clinical trials^21,22,50^, will potentially eliminate cell number constraints for cell therapy approaches to reverse diabetes. Much work is being done on scaling stem cell-derived insulin secreting cell therapy manufacturing, and our hydrogel injection molding macroencapsulation approach was designed to integrate with cell manufacturing pipelines to enable high-throughput production of encapsulated cell products, which could potentially lower costs. Of note, insulin-secreting stem cell sources do not have contaminating tissue like acinar in primary islet preparations, leading to more accurate macroencapsulation device cell loadings and potentially improved viability in vivo. Viacyte recently published clinical trial data^21,22^ using stem cell-derived insulin secreting cells in non-immune isolating devices transplanted subcutaneously, at a substantially higher density than is typical of 167 IEQ/µL (250,000 cells/µL); unsurprisingly, these devices demonstrate substantial central regions void of cellular material, likely due to cell death from hypoxia. Finally, while Viacyte found no evidence of teratoma formation^21^, stem cell derived cell sources raise potential safety risks, as evidenced by a recent aggressive teratoma formation in a diabetic patient receiving autologous induced pluripotent stem cell derived insulin secreting cells^51^, further supporting the need for fully retrievable macroencapsulation device approaches.

## 4 Conclusions

In this study, we establish that optimizing islet macroencapsulation device geometry can improve encapsulated cell survival and function over traditional geometries, facilitating safer cell therapy delivery to an isolated and retrievable site. We determined via computational modeling and in vitro validation that the optimal islet loading density of spiral macroencapsulation devices is 10 IEQ/µL, and an optimized device geometry reduces the required cell number for diabetes reversal or partial function in vivo in a syngeneic omentum transplant model. Altogether, toptimized device geometry greatly improves the clinical feasibility of macroencapsulated islet transplantation, and our hydrogel injection molding biomanufacturing method enhances the scalability of encapsulated cell therapy products.

## 5 Acknowledgements

This research was supported by the Juvenile Diabetes Research Foundation (grant 1-INO-2020-915-A-N), the Arizona Biomedical Research Centre New Investigator Award, and R01DK129858. The confocal microscope used in these studies was acquired by an NIH SIG award (1 S10 RR027154-01A1) and is housed in the Regenerative Medicine Imaging Facility at Arizona State University.

## 7 Author contributions

J.D.W. and A.E.E. developed the concept and design. J.D.W. and A.E.E. designed the study and analyzed data. A.E.E., Q.L., M.W.B., S.H., and S.R.B. conducted experiments and data collection, and J.D.W. assisted with data analysis. All authors contributed to writing the manuscript.

## 8 Data availability

The raw or processed data required to reproduce these findings are available upon request.

