## Supplemental Figures for "Hydrogel injection molded complex macroencapsulation device geometry improves long-term cell therapy viability and function in the rat omentum transplant site"

### Supplementary Information

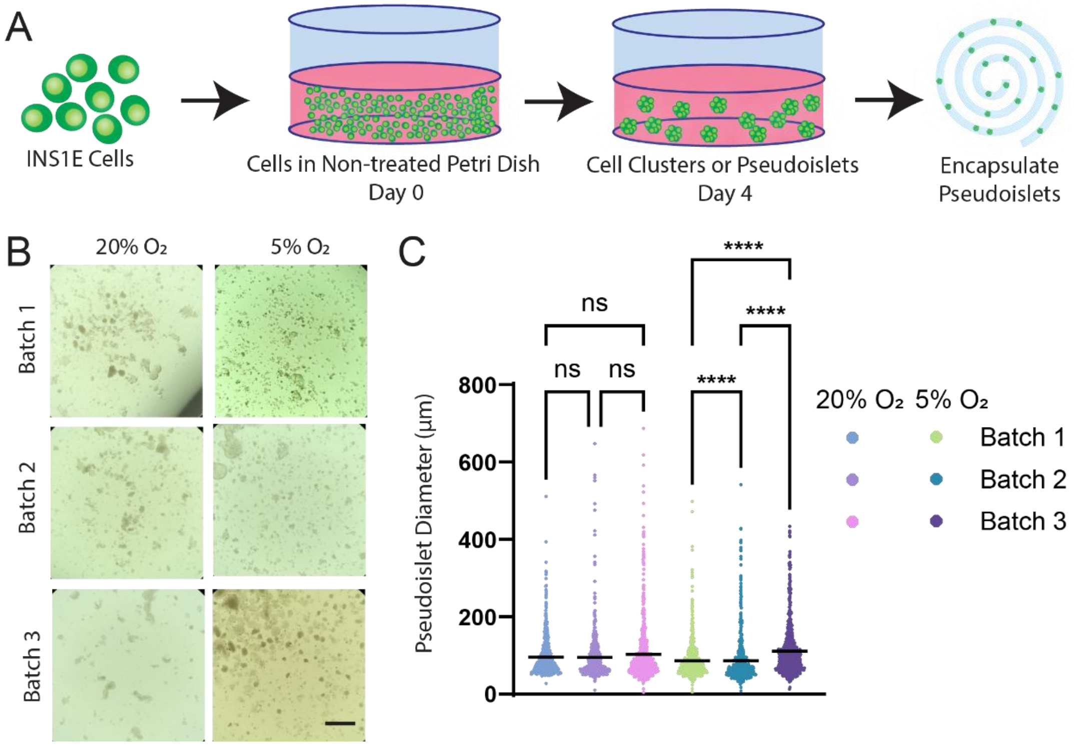


**Supplemental Figure 1**. **Fabrication and size analysis of INS-1 cell clusters or pseudoislets.** INS-1 cells were seeded in non-treated petri dishes of 100 mm diameter which were coated with 5 mL of anti-adherence rinsing solution. The INS-1 cells were cultured for 4 days in order to form pseudoislets (**A**). Images of the pseudoislets (**B**) were used to calculate the diameter distribution (**C**) of the pseudoislets. Pseudoislet diameters were analyzed by one-way ANOVA with Kruskal-Wallis multiple comparison test. ****P < 0.0001. Scale bars = 500 µm.

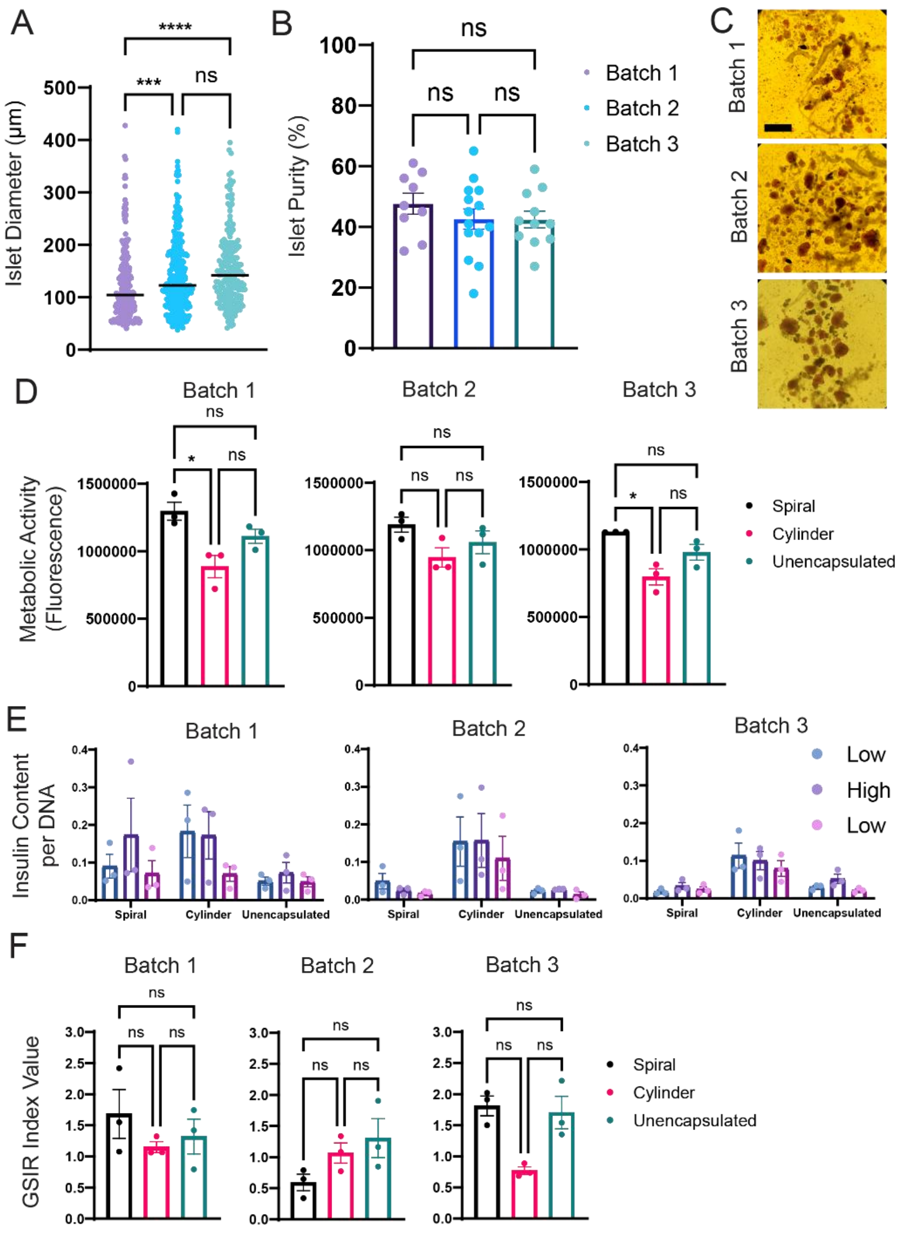


**Supplemental Figure 2. Effects of islet size distribution and purity on islet viability and function.** The islet size distribution (**A**) and islet purity (**B**) was evaluated prior to encapsulation from images (**C**) of islets stained with ditizone (DTZ). The individual batches of islets were investigated separately to determine the effects of islet size and purity on metabolic activity (**D**), insulin secretion (**E**), and GSIR index values (**F**). Each individual data point represents one hydrogel (n = 3 for each batch).Rat islet diameters and rat islet purity were analyzed by one-way ANOVA with Kruskal-Wallis multiple comparison test. Metabolic activity and GSIR index values analyzed by one-way ANOVA with Kruskal-Wallis multiple comparison test. *P < 0.05, ***P < 0.001, ****P < 0.0001. Error bars = SEM. Scale bars = 500 µm.

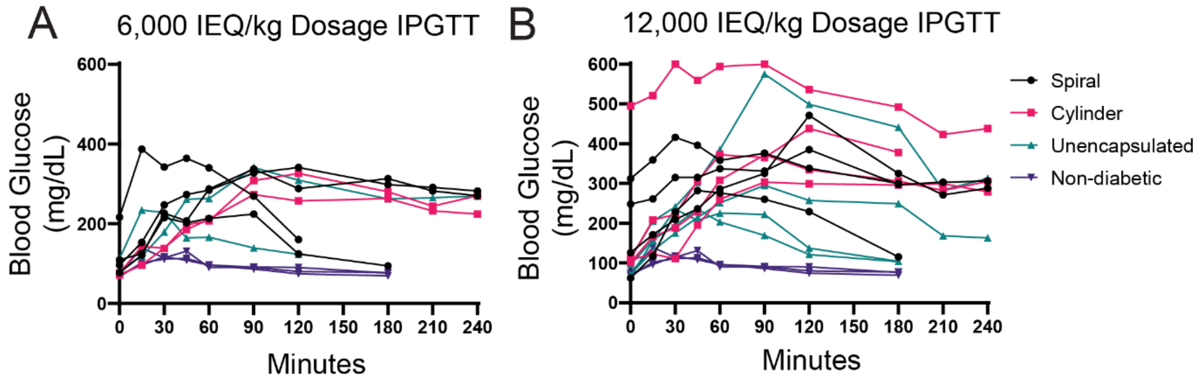


**Supplemental Figure 3. IPGTT values for individual rats.** The IPGTT test was performed on Week 14 for all groups in both the 6,000 IEQ/kg dosage study (**A**) and 12,000 IEQ/kg dosage study (**B**).

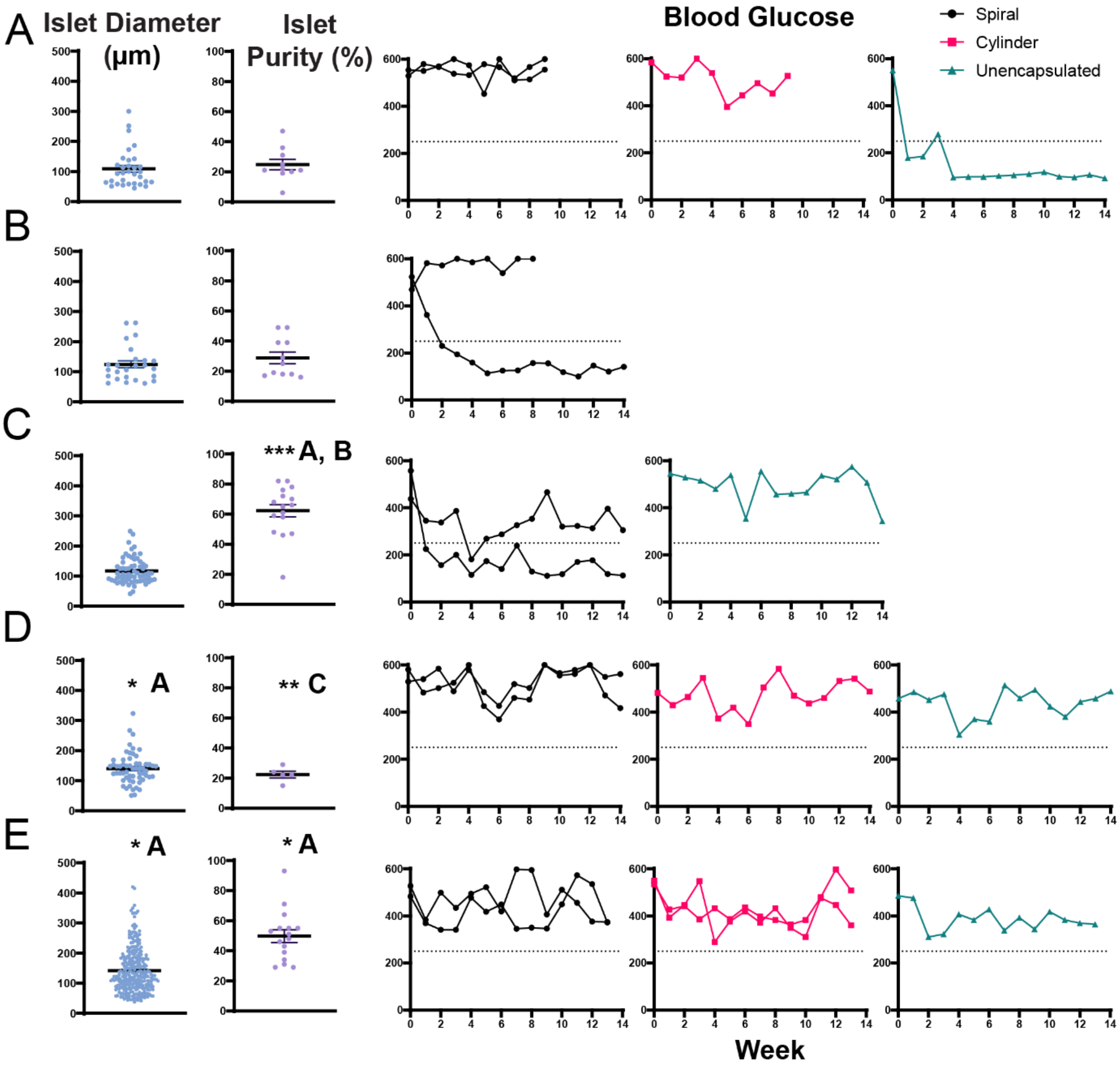


**Supplemental Figure 4. Effects of rat donor islet diameter and purity on individual transplants at a dosage of 6,000 IEQ/kg.** For each independent transplant, the islet diameter (µm) and islet purity (%) were calculated from images of DTZ-stained rat donor islets before encapsulation. Transplant 1 (**A**), Transplant 2 (**B**), Transplant 3 (**C**), Transplant 4 (**D**), and Transplant 5 (**E**). Islet diameter and purity was analyzed by one-way ANOVA with Kruskal-Wallis multiple comparison test. *P < 0.05, **P < 0.01, *** P < 0.001. Error bars = SEM.

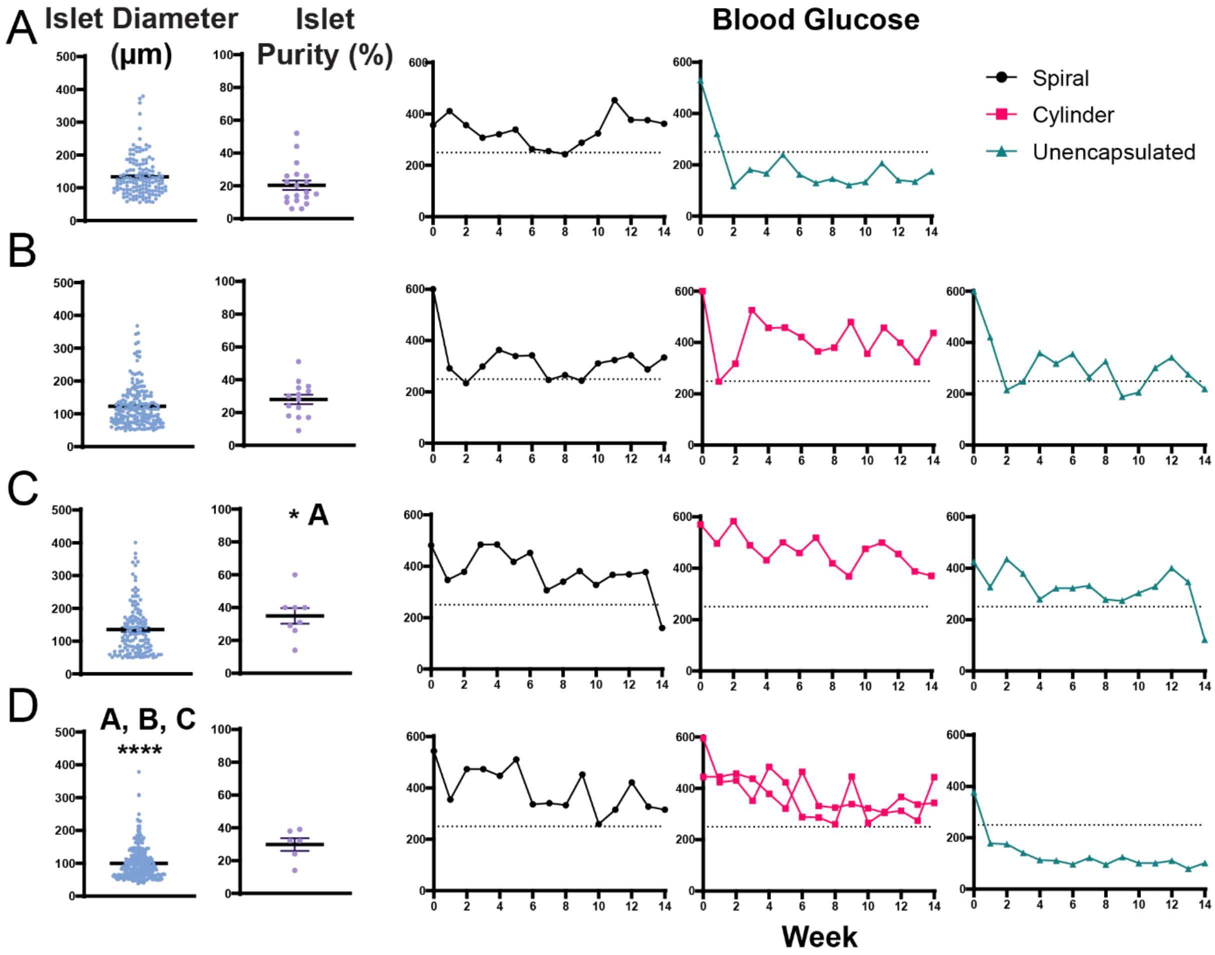


**Supplemental Figure 5. Effects of rat donor islet diameter and purity on individual transplants at a dosage of 12,000 IEQ/kg.** For each independent transplant, the islet diameter (µm) and islet purity (%) were calculated from images of DTZ-stained rat donor islets before encapsulation. Transplant 1 (**A**), Transplant 2 (**B**), Transplant 3 (**C**), and Transplant 4 (**D**). Islet diameter and purity was analyzed by one-way ANOVA with Kruskal-Wallis multiple comparison test. *P < 0.05, **** P < 0.0001. Error bars = SEM.

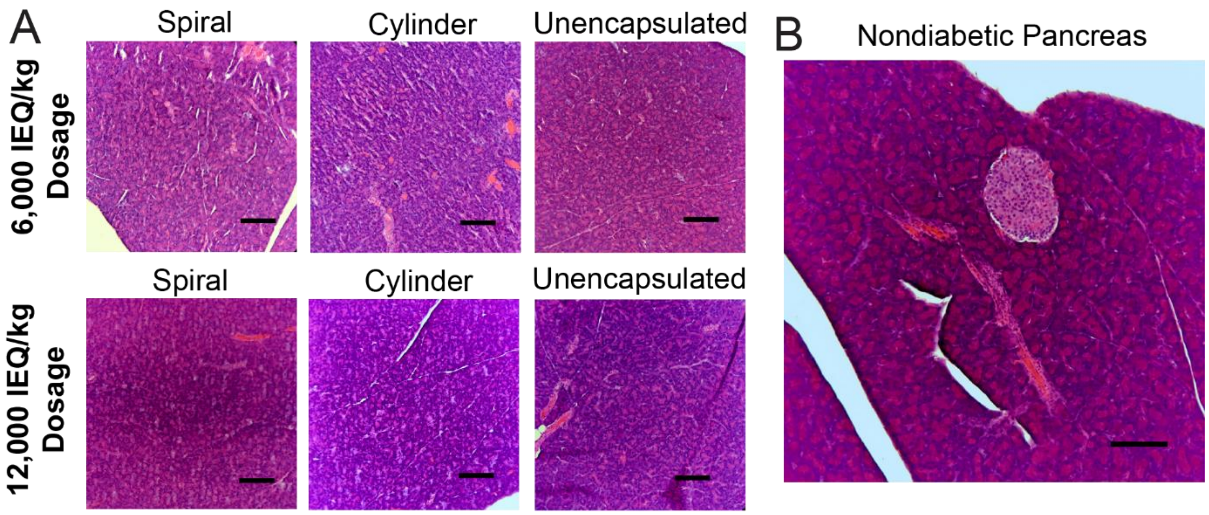


**Supplemental Figure 6. STZ pancreata analysis using H&E.** Sectioned tissue samples were stained and examined for functional islets (**A**). A healthy, non-diabetic pancreas was stained as a positive control of a functioning islet (**B**). Scale bar is 100 µm.
